## Supplementary file for "16S rRNA sequencing for profiling microbial communities in idiopathic granulomatous mastitis"

| **Item Number** | **Title** | **Main Text Reference** | **Supplementary Information Page Number** |
| --- | --- | --- | --- |
| Supplementary Table 1 | Idiopathic granulomatous mastitis (IGM) and lactational mastitis (LM) patients recruited, samples donated, and analysed. | Page 8  Page 11 | 3 |
| Supplementary Figure 1 | 16S rRNA gene map. | Page 9  Page 15 | 4 |
| Supplementary Table 2 | Median and inter-quartile range of alpha-diversity of breast pus and skin microbial populations from idiopathic granulomatous mastitis (IGM) and lactational mastitis (LM) patients. | Page 12  Page 13  Page 17 | 5 |
| Supplementary Figure 2 | Boxplot of the alpha-diversity of breast pus and skin microbial populations from idiopathic granulomatous mastitis (IGM) and lactational mastitis (LM) patients. Paired Wilcoxon sign ranked test comparing IGM-pus and IGM-skin samples. | Page 12 | 6-7 |
| Supplementary Figure 3 | Boxplot of the alpha-diversity of breast pus and skin microbial populations from idiopathic granulomatous mastitis (IGM) and lactational mastitis (LM) patients. Wilcoxon sign ranked test comparing (i) IGM-pus and LM-pus; and (ii) IGM-skin and LM-skin. | Page 12  Page 13 | 8-9 |
| Supplementary Figure 4 | Distribution of relative abundance of *Corynebacterium* genus in breast pus and skin samples from 3 lactational mastitis (LM) patients. | Page 13 | 10 |
| Supplementary Figure 5 | Relative abundance of *Corynebacterium kroppenstedtii* in breast **(a)** pus and **(b)** skin samples from 21 idiopathic granulomatous mastitis (IGM) and three controls with lactating mastitis (LM) (LM01, LM02, and LM03). | Page 13  Page 17 | 11-12 |
| Supplementary Table 3 | Statistically significant species (before and after adjustments, and after correcting for multiple comparisons) were identified from general linear models for determining multivariable association between sample type, covariates and microbial metagenomic features in paired pus and skin samples from idiopathic granulomatous mastitis (IGM) patients. | Page 17 | 13 |
| Supplementary Figure 6 | Scree plot of eigenvalues and multidimensional scaling (MDS) component numbers of the Bray-Curtis distance between breast pus and skin samples from Figure 3(a) and Figure 3(b). | Page 17 | 14 |
| Supplementary Table 4 | Statistically significant genera (after adjustments and correcting for multiple comparisons) were identified from general linear models for determining multivariable association between sample type, covariates and microbial metagenomic features in paired pus and skin samples from idiopathic granulomatous mastitis (IGM) patients. | Page 18 | 15 |

**Supplementary Table 1.** Idiopathic granulomatous mastitis (IGM) and lactational mastitis (LM) patients recruited, samples donated, and analysed. IGM: Idiopathic granulomatous mastitis; LM: lactational mastitis.

| Type of Mastitis | **IGM** | **LM** |
| --- | --- | --- |
| n | 26 | 6 |
| Skin Samples | 26 | 6 |
| Pus Samples | 21 | 4 |
| Paired Skin and Pus Samples | 21 | 4 |
| Metagenomic Analysis | 21 | 3 |

**Supplementary Figure 1.** 16S rRNA gene map. Variable regions are labelled from 1 to 9, in purple blocks, and conserved regions are in grey. The entire gene is approximately 1,600 base pairs long. Yellow arrows: Forward and reverse primers used in Yu, et al., 2016; Blue arrows: Forward and reverse primers used in our study. F: Forward primer; R: Reverse primer; V: 16S rRNA gene variable region; bp: Base pairs.

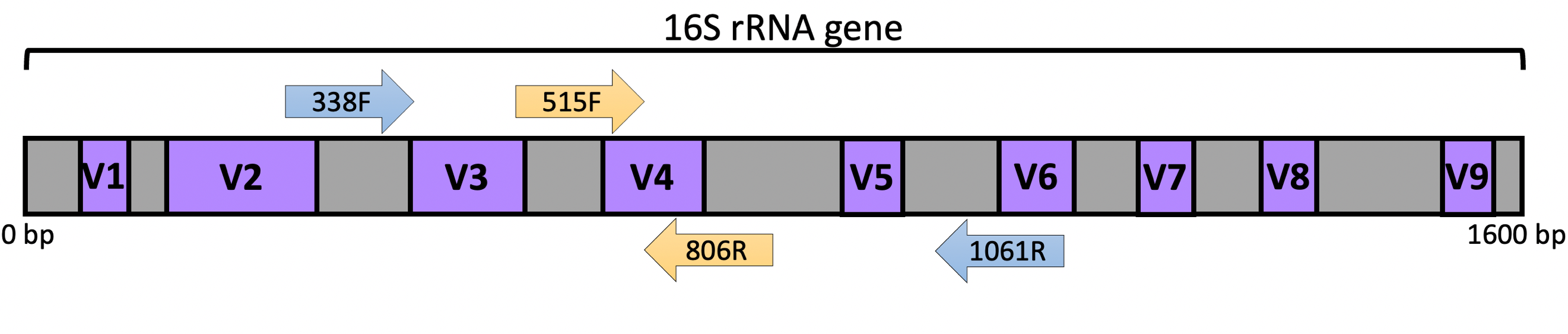

**Supplementary Table 2.** Median and inter-quartile range of alpha-diversity of breast pus and skin microbial populations from idiopathic granulomatous mastitis (IGM) and lactational mastitis (LM) patients, using Shannon and Simpson indices. Formula used for indices calculation can be found in Supplementary Formula 1. Shannon index is an information statistic index that measures the number of different genera, and the evenness in distribution of the genera. More unique genera, or more genera evenness increases Shannon index. Simpson index is a dominance index that gives more weight to common and dominant species. Rare species with low abundance is less important in this diversity index. IGM: Idiopathic granulomatous mastitis; LM: lactational mastitis.

|  | **Shannon** | | | | **Simpson** | | | |
| --- | --- | --- | --- | --- | --- | --- | --- | --- |
|  | **IGM** | | **LM** | | **IGM** | | **LM** | |
|  | **Pus** | **Skin** | **Pus** | **Skin** | **Pus** | **Skin** | **Pus** | **Skin** |
| 25% | 1.91 | 1.46 | 1.07 | 1.07 | 0.75 | 0.54 | 0.45 | 0.45 |
| median | 2.65 | 1.80 | 1.66 | 1.66 | 0.88 | 0.74 | 0.73 | 0.73 |
| 75% | 2.92 | 2.36 | 2.26 | 2.26 | 0.92 | 0.82 | 0.81 | 0.81 |

**Supplementary Figure 2.** Boxplot of the alpha-diversity of breast pus and skin microbial populations from idiopathic granulomatous mastitis (IGM) and lactational mastitis (LM) patients, using **(a)** Shannon and **(b)** Simpson indices. Formula used for indices calculation can be found in Supplementary Formula 1.

**(a)** Shannon index is an information statistic index that measures the number of different genera, and the evenness in distribution of the genera. More unique genera, or more genera evenness increases Shannon index.

**(b)** Simpson index is a dominance index that gives more weight to common and dominant species. Rare species with low abundance is less important in this diversity index.

Paired Wilcoxon sign ranked test found significant higher diversity in IGM pus samples, compared to the corresponding paired IGM skin sample, for both Shannon (p=0.022) and Simpson (p=0.07) indices. IGM: Idiopathic granulomatous mastitis; LM: lactational mastitis.

**Supplementary Figure 2.**

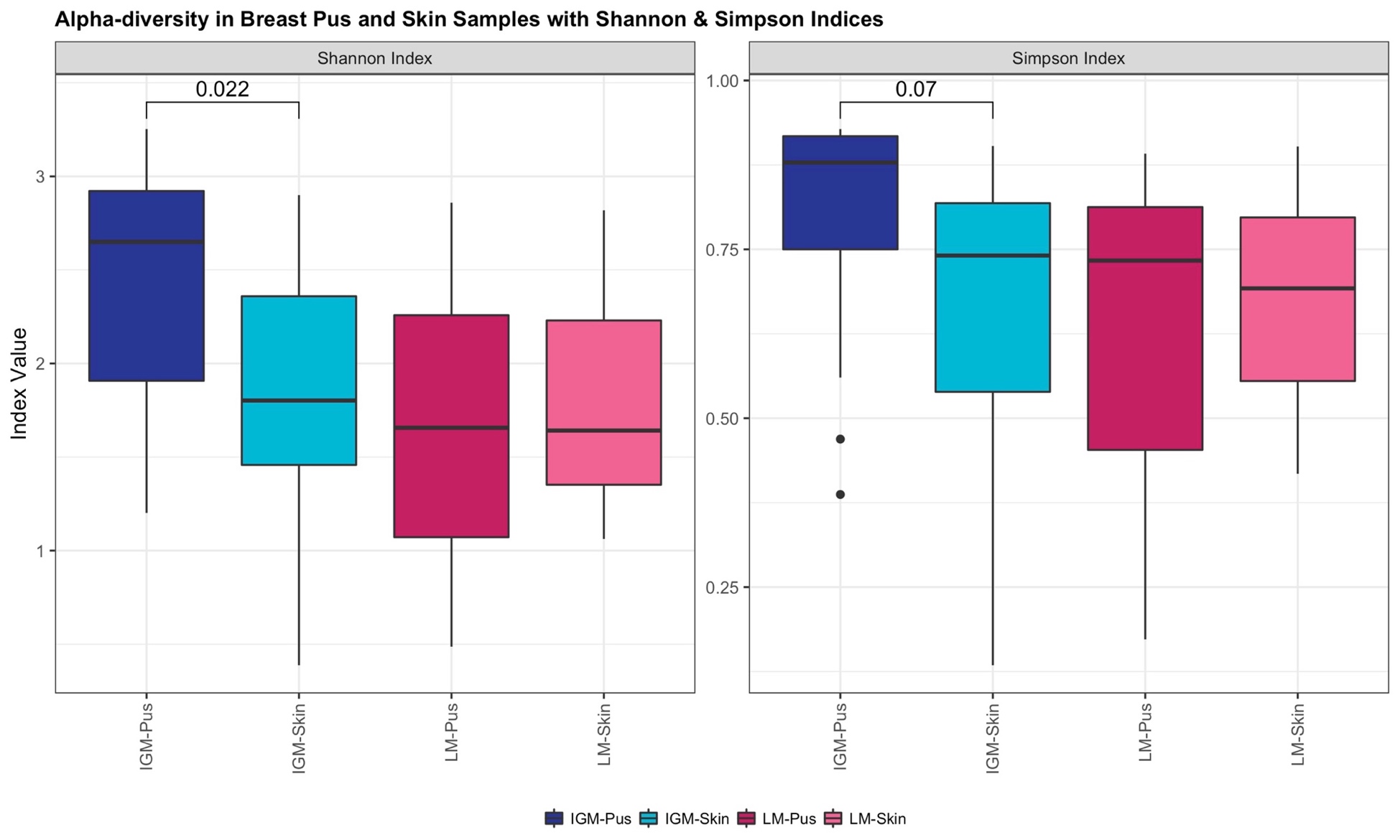

**Supplementary Figure 3.** Boxplot of the alpha-diversity of breast pus and skin microbial populations from idiopathic granulomatous mastitis (IGM) and lactational mastitis (LM) patients, using **(a)** Shannon and **(b)** Simpson indices. Formula used for indices calculation can be found in Supplementary Formula 1.

(a) Shannon index is an information statistic index that measures the number of different genera, and the evenness in distribution of the genera. More unique genera, or more genera evenness increases Shannon index.

(b) Simpson index is a dominance index that gives more weight to common and dominant species. Rare species with low abundance is less important in this diversity index.

Wilcoxon sign ranked test did not find significant differences between IGM Pus and LM Pus, or IGM Skin and LM Skin samples, in either indices. IGM: Idiopathic granulomatous mastitis; LM: lactational mastitis.

**Supplementary Figure 3.**

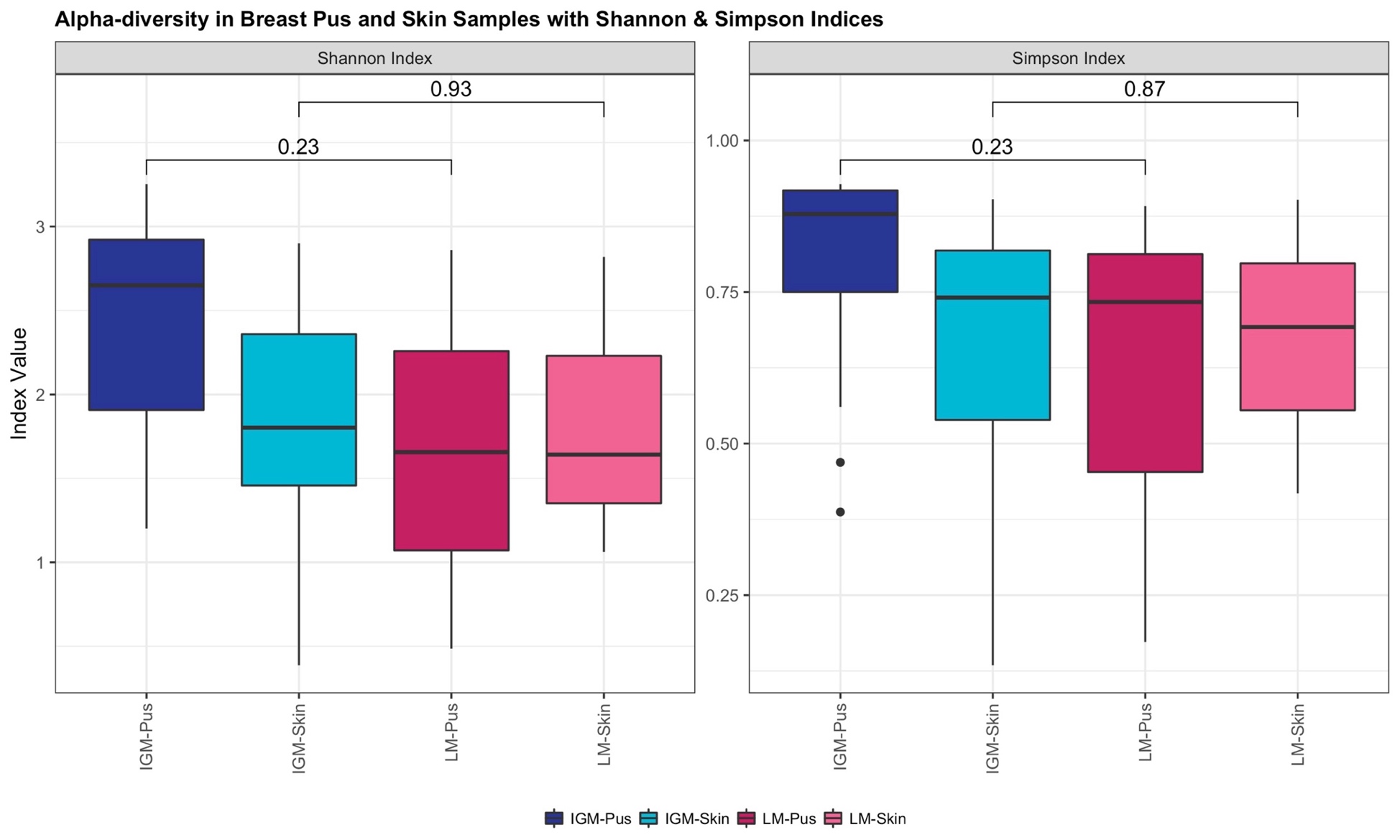

**Supplementary Figure 4.** Distribution of relative abundance of *Corynebacterium* genus in breast pus and skin samples from 3 lactational mastitis (LM) patients. Median *Corynebacterium* relative abundance in LM pus samples is 1.4% (interquartile range = 0.9-5.1%), compared to 5.8% (interquartile range = 5.6-7.9%) in LM skin samples. Paired Wilcoxon sign ranked test found no significant difference in *Corynebacterium* relative abundance between paired LM skin and pus samples (p = 0.25). LM: lactational mastitis; p: p-value; Wilcoxon: Wilcoxon paired sign ranked test.

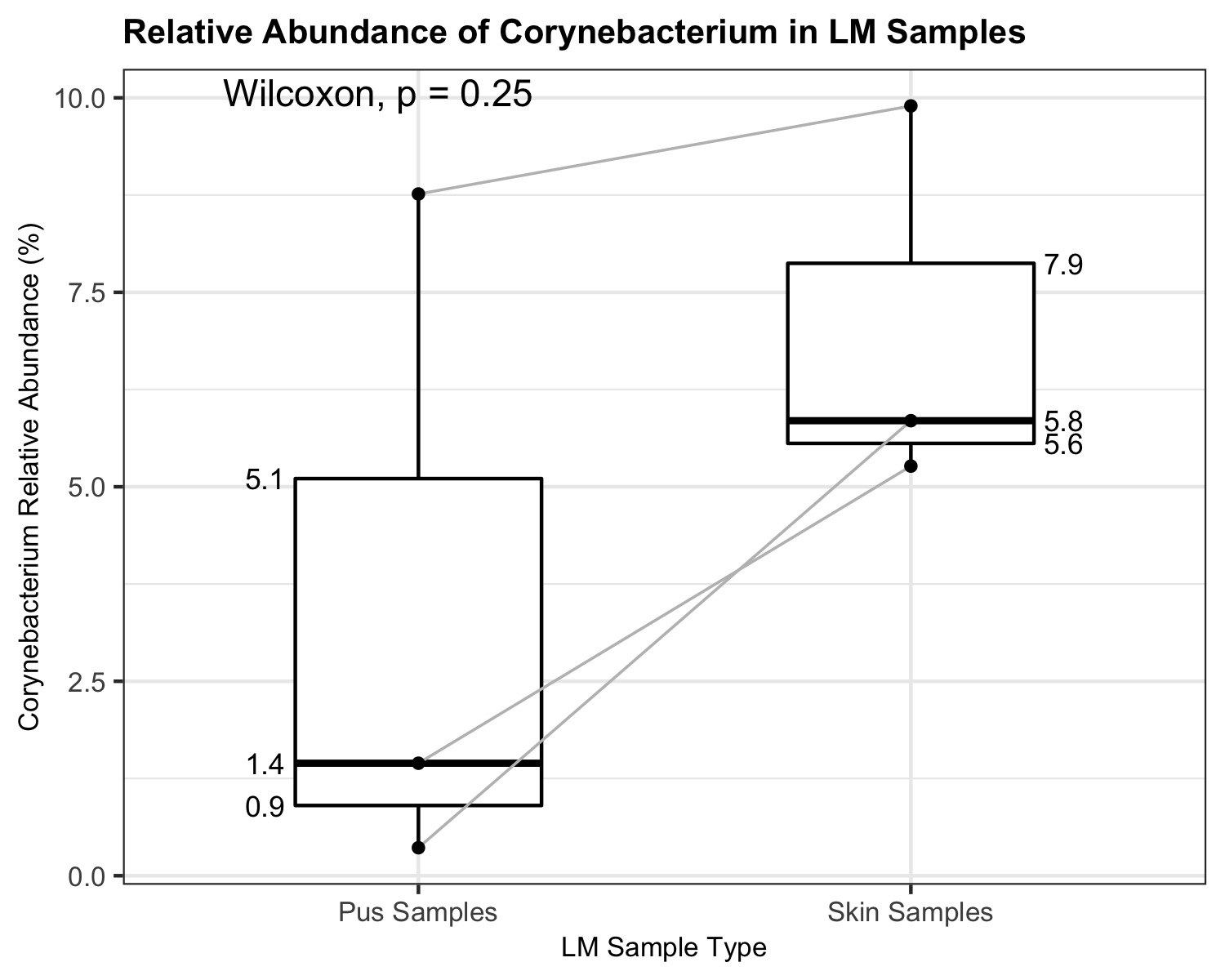

**Supplementary Figure 5.** Relative abundance of *Corynebacterium kroppenstedtii* in breast **(a)** pus and **(b)** skin samples from 21 idiopathic granulomatous mastitis (IGM) and three controls with lactating mastitis (LM) (LM01, LM02, and LM03). The left-end bars in both panels represent the IGM patients, and the right-end bars in both panels represent the LM patients, with corresponding colours indicating mastitis type shown in the bottom-left shared legend for **(a)** and **(b)**. The percentage of *Corynebacterium kroppenstedtii* relative abundance is indicated above each bar, presented to 1 decimal place. Duration of antibiotic treatment prior to sample collection are presented as symbols above each bar, as four categories: Less than 2 weeks before sample collection, more than 2 weeks before sample collection, missing duration, and no antibiotic treatment. The corresponding symbols are indicated in the bottom-left shared legend for **(a)** and **(b)**.

**(a)** For the left panel of **pus** samples, the patients are arranged in decreasing relative abundance of *Corynebacterium kroppenstedtii* in **pus** samples within mastitis type.

**(b)** For the right panel of **skin** samples, the patients are arranged following patient order in **(a)** i.e. decreasing relative abundance of *Corynebacterium kroppenstedtii* in **pus** samples within mastitis type.

Distribution of relative abundance of *Corynebacterium kroppenstedtii* in breast pus and skin samples from 21 IGM patients is also displayed in **(a)**. Median *Corynebacterium* relative abundance in IGM pus samples is 2.1% (interquartile range = 0.3-7.0%), compared to 0.1% (interquartile range = 0.0-0.9%) in IGM skin samples. Paired Wilcoxon sign ranked test was significantly higher for *Corynebacterium kroppenstedtii* relative abundance in IGM pus samples than the paired skin samples (p = 0.022).

IGM: Idiopathic granulomatous mastitis; LM: lactational mastitis; p: p-value; Wilcoxon: Wilcoxon paired sign ranked test.

**Supplementary Figure 5.**

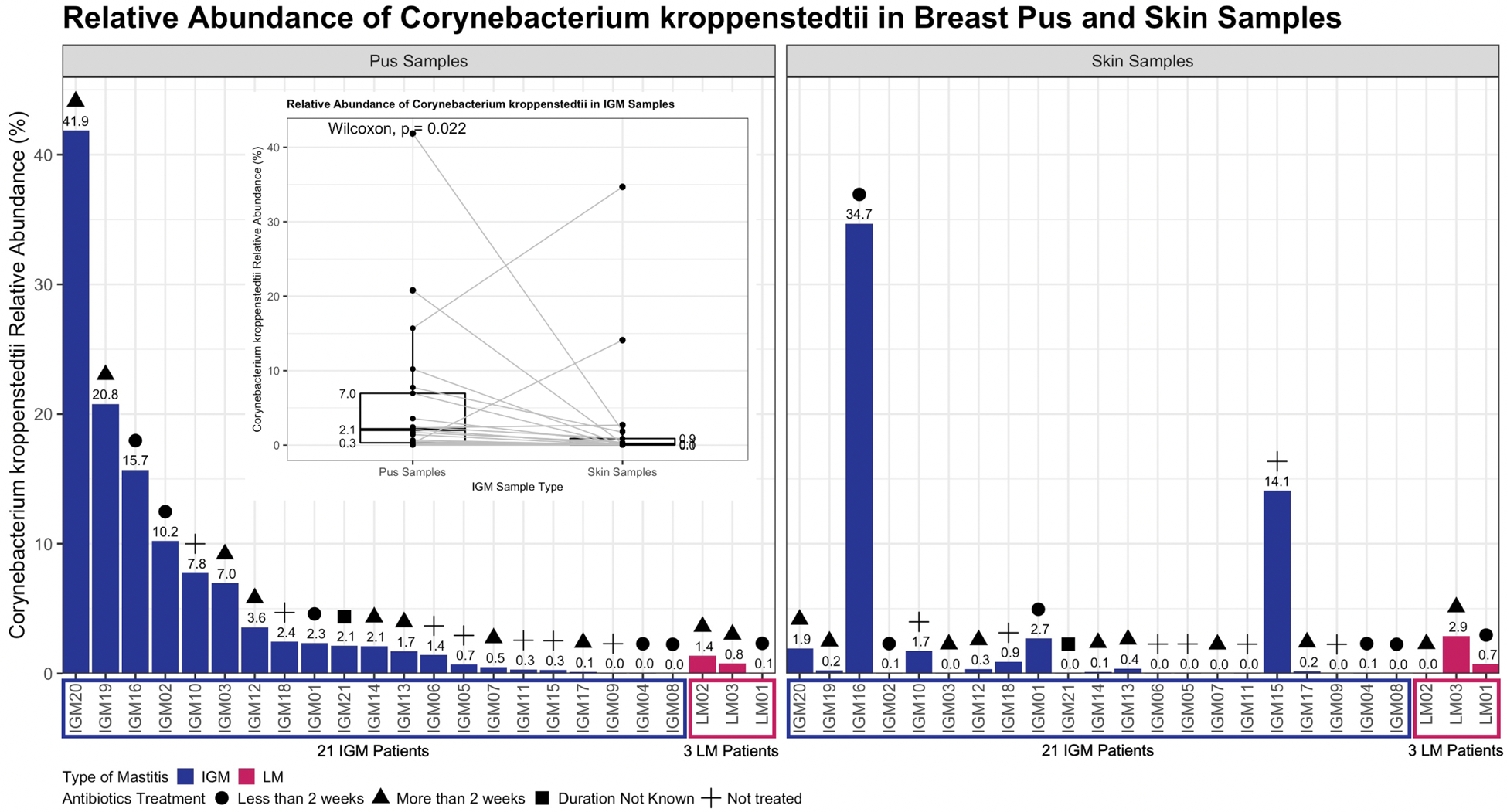

**Supplementary Table 3.** Statistically significant species (before and after adjustments, and after correcting for multiple comparisons) were identified from general linear models for determining multivariable association between sample type, covariates and microbial metagenomic features in paired pus and skin samples from idiopathic granulomatous mastitis (IGM) patients. Beta-estimates, standard deviations, crude and adjusted p-values, and crude and adjusted q-values are reported in Table 4. The median relative abundance (%) and the interquartile range (IQR) for IGM pus and skin samples, and lactational mastitis (LM) pus and skin samples are presented. IQR: Interquartile range; IGM: Idiopathic granulomatous mastitis; LM: lactational mastitis.

| **Species** | **Relative abundance (%) (IQR, %)** | | | |
| --- | --- | --- | --- | --- |
|  | **IGM** | | **LM** | |
|  | Pus | Skin | Pus | Skin |
| *Acinetobacter schindleri* | 0.94 (0.04-1.72) | 0.00 (0.00-0.00) | 0.00 (0.00-0.88) | 0.00 (0.00-0.00) |
| *Rothia mucilaginosa* | 0.27 (0.12-0.53) | 0.00 (0.00-0.02) | 0.02 (0.01-0.28) | 0.00 (0.00-7.12) |
| *Lactobacillus iners* | 0.00 (0.00-0.00) | 0.00 (0.00-0.01) | 0.00 (0.00-0.00) | 0.00 (0.00-0.00) |
| *Corynebacterium kroppenstedtii* | 2.10 (0.31-6.96) | 0.12 (0.02-0.89) | 0.76 (0.42-1.07) | 0.72 (0.38-1.79) |
| *Roseomonas mucosa* | 0.00 (0.00-0.00) | 0.03 (0.00-0.13) | 0.00 (0.00-0.00) | 0.01 (0.00-0.38) |
| *Kocuria rhizophila* | 0.00 (0.00-0.00) | 0.00 (0.00-0.06) | 0.00 (0.00-0.00) | 0.00 (0.00-0.00) |

**Supplementary Figure 6.** Scree plot of eigenvalues and multidimensional scaling (MDS) component numbers of the Bray-Curtis distance between breast pus and skin samples from Figure 3(a) and Figure 3(b).

**(a)** Eigenvalues for Figure 3(a) MDS of Bray-Curtis distance between breast pus and skin samples for idiopathic granulomatous mastitis (IGM) and lactational mastitis (LM) patients.

**(b)** Eigenvalues for Figure 3(b) MDS of Bray-Curtis distance between breast pus and skin samples for IGM patients only.

IGM: Idiopathic granulomatous mastitis; LM: lactational mastitis.

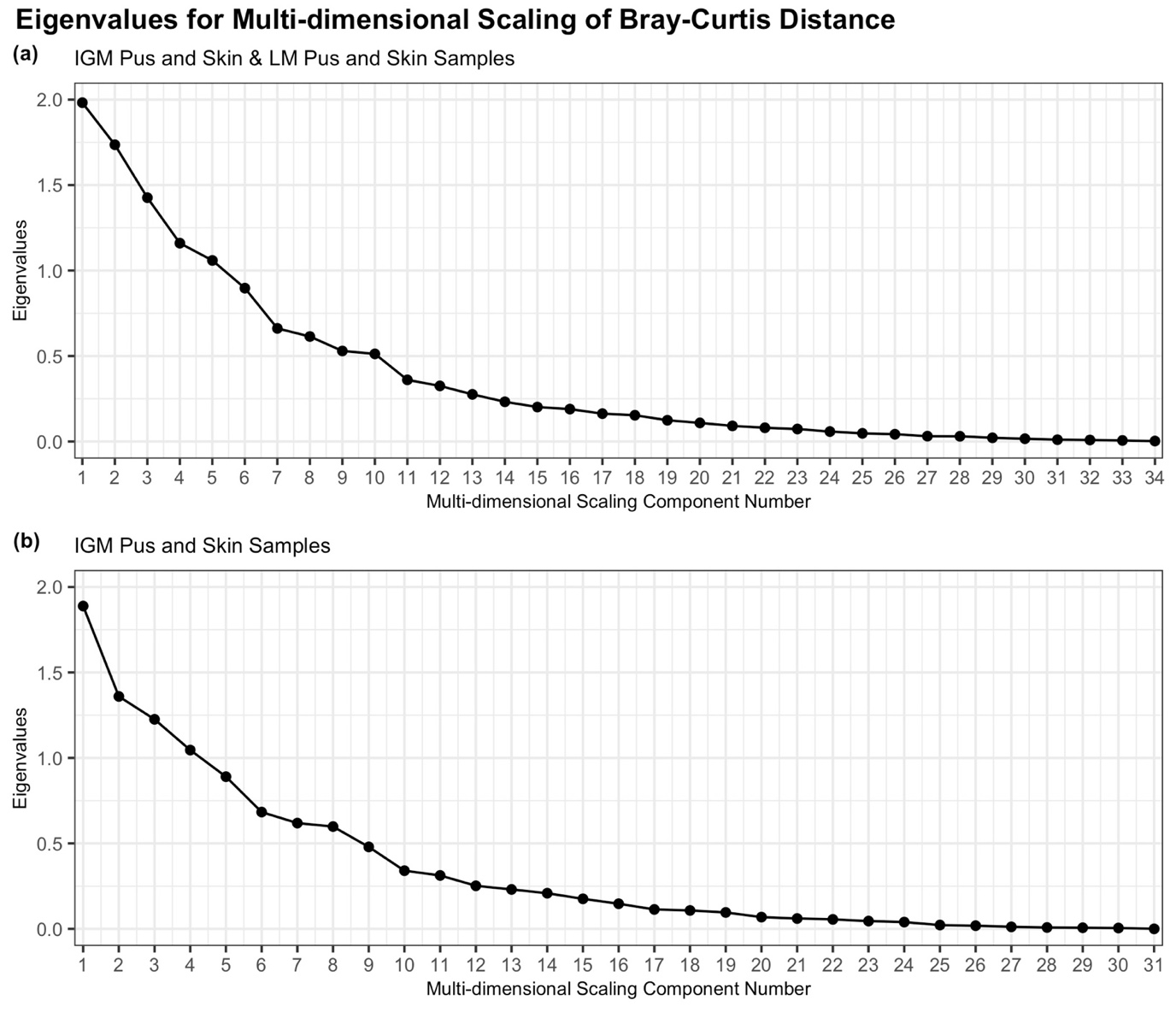

**Supplementary Table 4.** Statistically significant genera (after adjustments and correcting for multiple comparisons) were identified from general linear models for determining multivariable association between sample type, covariates and microbial metagenomic features in paired pus and skin samples from idiopathic granulomatous mastitis (IGM) patients. Beta-estimates, standard deviations, crude and adjusted p-values, and crude and adjusted q-values are reported in Table 3. The median relative abundance (%) and the interquartile range (IQR) for IGM pus and skin samples, and lactational mastitis (LM) pus and skin samples are presented. IQR: Interquartile range; IGM: Idiopathic granulomatous mastitis; LM: lactational mastitis.

| **Genus** | **Relative abundance (%) (IQR, %)** | | | |
| --- | --- | --- | --- | --- |
|  | **IGM** | | **LM** | |
|  | Pus | Skin | Pus | Skin |
| *Ochrobactrum* | 1.75 (1.29-2.81) | 0.00 (0.00-0.00) | 0.09 (0.04-0.78) | 0.00 (0.00-0.00) |
| *Delftia* | 4.47 (1.85-10.01) | 0.01 (0.01-0.01) | 0.26 (0.14-2.17) | 0.01 (0.01-0.01) |
| *Anaerobacillus* | 0.60 (0.22-0.98) | 0.00 (0.00-0.00) | 0.00 (0.00-0.02) | 0.00 (0.00-0.00) |
| *Gordonia* | 0.69 (0.32-1.06) | 0.02 (0.01-0.05) | 0.05 (0.03-0.55) | 0.08 (0.04-0.18) |
| *Methylobacterium* | 0.00 (0.00-0.00) | 0.02 (0.00-0.04) | 0.00 (0.00-0.00) | 0.01 (0.01-0.01) |
| *Fusobacterium* | 0.92 (0.35-1.43) | 0.01 (0.00-0.03) | 0.03 (0.03-1.53) | 0.01 (0.01-0.02) |
| *Sphingobium* | 0.00 (0.00-0.00) | 0.00 (0.00-0.02) | 0.00 (0.00-0.08) | 0.00 (0.00-0.00) |
| *Alkanindiges* | 0.29 (0.02-0.43) | 0.00 (0.00-0.01) | 0.02 (0.01-0.19) | 0.00 (0.00-0.00) |
| *Streptococcus* | 3.51 (1.76-5.09) | 0.23 (0.14-0.58) | 3.95 (2.25-5.69) | 8.94 (7.06-29.98) |
| *Achromobacter* | 0.00 (0.00-0.00) | 0.02 (0.00-0.10) | 0.00 (0.00-0.00) | 0.02 (0.01-0.03) |
| *Capnocytophaga* | 0.00 (0.00-0.00) | 0.00 (0.00-0.03) | 0.00 (0.00-0.00) | 0.01 (0.01-0.01) |
| *Mycobacterium* | 0.00 (0.00-0.00) | 0.04 (0.01-0.08) | 0.00 (0.00-0.00) | 0.08 (0.05-0.18) |
| *Novosphingobium* | 0.00 (0.00-0.00) | 0.01 (0.00-0.01) | 0.00 (0.00-0.00) | 0.01 (0.01-0.07) |
| *Peptoniphilus* | 0.45 (0.31-0.68) | 0.02 (0.01-0.11) | 0.02 (0.01-0.33) | 0.01 (0.00-0.01) |
| *Rothia* | 0.29 (0.12-0.53) | 0.02 (0.00-0.03) | 0.02 (0.01-0.28) | 0.03 (0.02-7.13) |
| *Finegoldia* | 0.94 (0.39-1.13) | 0.07 (0.02-0.10) | 0.03 (0.02-0.62) | 0.01 (0.01-0.02) |
| *Burkholderia* | 0.00 (0.00-0.00) | 0.01 (0.00-0.04) | 0.00 (0.00-0.11) | 0.00 (0.00-0.01) |
| *Roseomonas* | 0.00 (0.00-0.00) | 0.03 (0.00-0.13) | 0.00 (0.00-0.00) | 0.01 (0.00-0.38) |
| *Anaerococcus* | 0.69 (0.40-0.92) | 0.06 (0.02-0.19) | 0.17 (0.10-0.48) | 0.05 (0.02-0.13) |
| *Agrobacterium* | 0.00 (0.00-0.00) | 0.01 (0.01-0.07) | 0.05 (0.03-0.06) | 0.01 (0.00-0.04) |
| *Hydrogenophaga* | 0.00 (0.00-0.00) | 0.00 (0.00-0.02) | 0.00 (0.00-0.00) | 0.00 (0.00-0.00) |
| *Peptostreptococcus* | 0.00 (0.00-0.00) | 0.00 (0.00-0.01) | 0.00 (0.00-0.00) | 0.00 (0.00-0.00) |
| *Dermabacter* | 0.00 (0.00-0.00) | 0.03 (0.00-0.07) | 0.00 (0.00-0.00) | 0.05 (0.03-0.08) |
| *Kocuria* | 0.00 (0.00-0.00) | 0.03 (0.00-0.07) | 0.00 (0.00-0.00) | 0.00 (0.00-0.00) |
